# Comparing dopamine release, uptake, and D2 autoreceptor function across the ventromedial to dorsolateral striatum in adolescent and adult rats

**DOI:** 10.1101/806638

**Authors:** Elizabeth G. Pitts, Taylor A. Stowe, Brooke Christensen, Mark J. Ferris

**Affiliations:** Department of Physiology and Pharmacology, Wake Forest School of Medicine, Winston-Salem, North Carolina 27157

**Keywords:** Adolescence, Dopamine, Striatum, Voltammetry

## Abstract

Adolescents have increased vulnerability for the development of a range of psychiatric disorders, including substance abuse disorders (SUD) and mood disorders. Adolescents also have increased rates of sensation seeking and risk taking. The mesolimbic dopamine system plays a role in all these behaviors and disorders and reorganization of the dopamine system during adolescence may be important in mediating these developmental changes in behavior and vulnerability. Here, we used *ex vivo* fast scan cyclic voltammetry to examine developmental differences in dopamine release and its local circuitry regulation across the striatum. We found that adolescents have significantly decreased dopamine release in the nucleus accumbens core across a range of stimulation frequencies that model tonic and phasic firing of mesolimbic dopamine neurons. We show this is not mediated by differences in rate of dopamine uptake, but may be driven by hypersensitive dopamine autoreceptors, indicated by increased inhibition in dopamine release following agonism of D2/D3 receptors, in the adolescent nucleus accumbens core. Additionally, we observed increases in dopamine uptake in the dorsomedial striatum. No other significant differences between release, uptake, or D2 autoreceptor function was observed between adolescent and adult rats in all brain areas tested (nucleus accumbens shell, nucleus accumbens core, dorsomedial striatum, and dorsolateral striatum). These developmental differences in dopamine release may be important in mediating some of the unique behavioral repertoire seen in adolescents, such as increases in sensation seeking, and its associated vulnerabilities.

## 1. Introduction

Adolescence is a developmental time-period characterized by increases in reward sensitivity, sensation seeking, and peer-focused social behavior (Spear, 2000, 2011, 2013; Nelson et al., 2005). It has been proposed that these changes in behavior are biologically advantageous, supporting the development of independence necessary for the movement from childhood to adulthood (Spear, 2000). However, this is also a time of increased vulnerability to both the consequences of increased risk-taking and the development of a range of psychiatric disorders, including depression, schizophrenia, and substance use disorders (SUDs) (Casey et al., 2008; Fareri et al., 2008; Steinberg, 2008).

Developmental reorganization of the mesolimbic dopamine system is hypothesized to drive altered reward-related decision making, including increasing risk-taking behaviors, and to play a role in the increased vulnerability to a variety of psychiatric disorders, including SUDs and mood and anxiety disorders, that is seen during adolescence (Spear, 2013; Nelson et al., 2005; Wahlstrom et al., 2010a,b). A number of changes occur to the mesolimbic dopamine system during adolescence (see Wahlstrom et al., 2010a,b; Padmanabhan & Luna, 2014). For example, D1 and D2 receptor density peaks in the striatum during adolescence (Teicher et al., 1995; Tarazi et al., 1999; Philpot et al., 2009), as does dopamine neuron firing rates (McCutcheon et al., 2012).

Despite the importance of changes to the dopamine system on adolescent behaviors and vulnerabilities, changes in striatal dopamine levels and their regulation are not well understood. Previous studies have used microdialysis to compare extracellular dopamine levels in the ventral striatum of adolescents and adults, but these studies have found conflicting results showing both increased and decreased levels of dopamine in adolescent animals (Badanich et al., 2006; Cao et al., 2007; Philpot et al., 2009). Additionally, one study examined rapid dopamine release and uptake in the striatum using anesthetized *in vivo* fast scan cyclic voltammetry (FSCV) and found decreased dopamine release in the striatum of adolescents (Stamford, 1989). However, no one has yet compared rapid dopamine release and its modulation in adolescent and adult rats throughout the striatum. *Ex vivo* FSCV provides unique advantages for studying local circuitry modulation of dopamine release. It has high spatial and temporal resolution and can be used to examine uptake kinetics and the role of transporters and autoreceptors in mediating dopamine release (Ferris et al., 2013). Additionally, electrical stimulation can model both tonic (∼ 4-5 Hz) and phasic (2-5 spikes at 20-100 Hz) neuronal firing, which may be particularly important for the salience of rewarding information in adolescence (Luciana et al., 2012).

Here, we used *ex vivo* FSCV to compare sub-second dopamine release and uptake in adults and adolescents across several regions of the striatum that may play a role in mediating behaviors and vulnerability unique to adolescence. In particular, we focused on the nucleus accumbens shell (NAc shell) and core (NAc core), which are important in incentive motivated behavior and reward-related learning (see Di Chiara, 2002; Saddoris et al., 2013). Additionally, the NAc core plays a role in goal-directed decision making (Saddoris et al., 2013). We also examined the dorsomedial striatum (DMS) and the dorsolateral striatum (DLS), which are necessary for flexible, goal-directed actions and habit formation, respectively (see Balleine & O’Doherty, 2010). Various stimulation parameters were used to model a range of dopamine neuron firing patterns. We then used selective D2/D3 agonists and antagonists to examine the potential role of autoreceptors in mediating developmental changes in dopamine release. We found developmental differences in dopamine release and its local circuitry regulation, particularly in the NAc core.

## 2. Materials and Methods

### 2.1 Animals

Adolescent (P30-35) and adult (P70-90) male Sprague-Dawley rats (Envigo, Huntingdon, UK) were maintained on a 12:12 h reverse light/dark cycle (4:00 a.m. lights off; 4:00 p.m. lights on) with food and water available *ad libitum.* All animals were maintained according to the National Institutes of Health guidelines in Association for Assessment and Accreditation of Laboratory Animal Care accredited facilities. All experimental protocols were approved by the Institutional Animal Care and Use Committee at Wake Forest School of Medicine.

### 2.2 Slice preparation

Rats were anesthetized with isoflurane and euthanized by decapitation in a ventilated area. Brains were rapidly removed and transferred into pre-oxygenated (95% O_2_ / 5% CO_2_) artificial cerebral spinal fluid (aCSF) containing (in mM): NaCl (126), KCl (2.5), monobasic NaH_2_PO_4_ (1.2), CaCl_2_ (2.4), MgCl_2_ (1.2), NaHCO_3_ (25), dextrose (D-glucose) (11), and L-ascorbic acid (0.4). Tissue was sectioned into 400 μm-thick coronal slices on a compresstome^®^ VF-300 vibrating microtome (Precisionary Instruments Inc., San Jose, CA). Brain slices were transferred to testing chambers containing oxygenated aCSF (32 °C) flowing at 1 mL/min.

### 2.3 Ex vivo voltammetry

*Ex vivo* FSCV was used to characterize presynaptic dopamine release in the NA shell, NAc, DMS, and DLS (Figure 1A). A carbon-fiber recording microelectrode (100–200 μM length, 7 μM diameter) was placed 100-150 μM from a bipolar stimulating electrode. Extracellular dopamine was recorded by applying a triangular waveform from −0.4 to 1.2 V and back to −0.4 (Ag vs AgCl) at a scan rate of 400 V/s.

**Figure 1.**
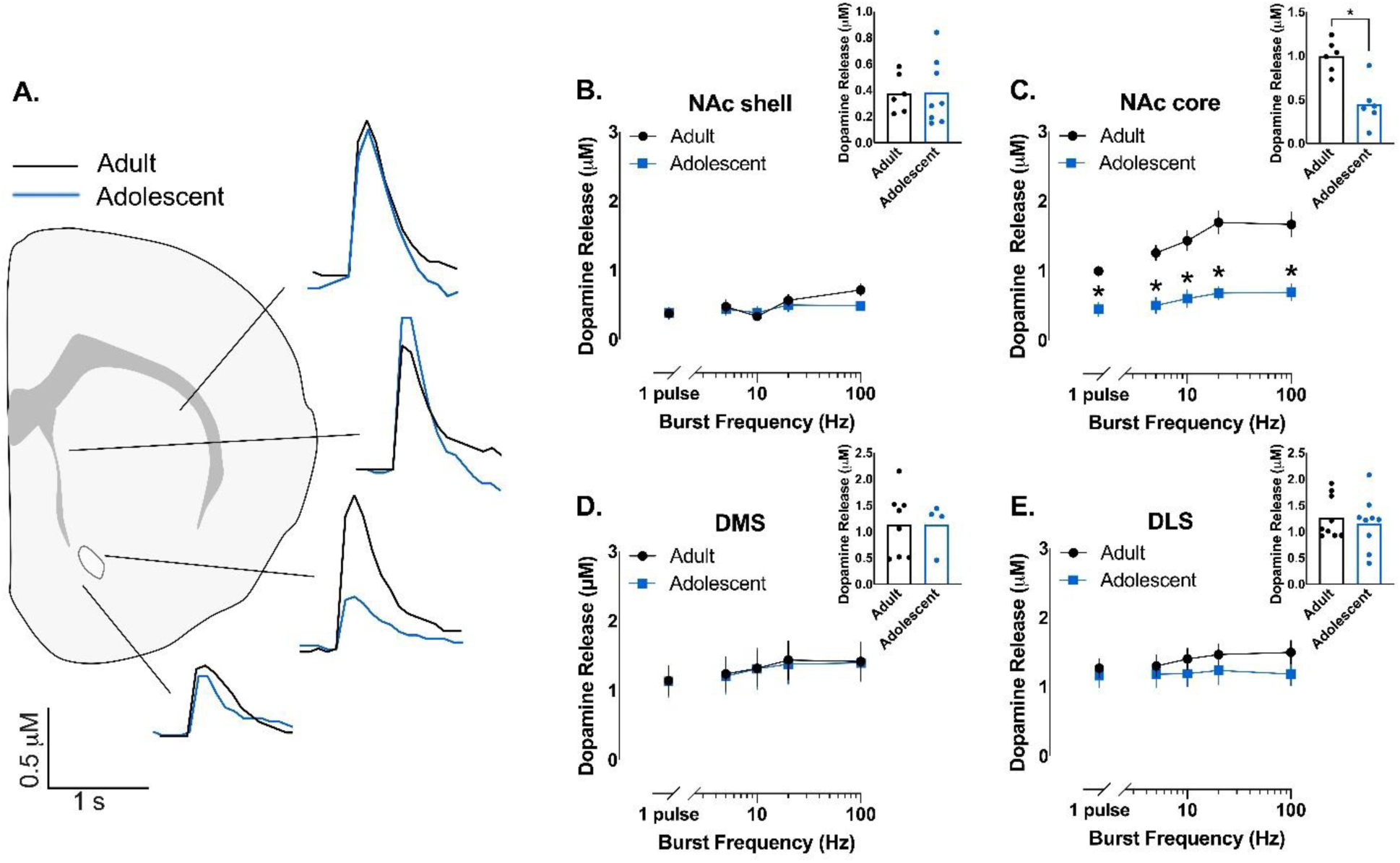
Adolescents have decreased dopamine release in the nucleus accumbens core. (A) *Ex vivo* fast scan cyclic voltammetry was used to examine dopamine release throughout the striatum in adult and adolescent male rats. Representative traces of stimulated single-pulse dopamine release in the nucleus accumbens shell (NA shell), nucleus accumbens core (NAc core), dorsomedial striatum (DMS), and dorsolateral striatum (DLS) are shown. (B) Dopamine release in the NA shell was not different in adult and adolescent animals. (C) However, dopamine release was significantly lower in adolescent than adult rats in the NAc core across a range of stimulation parameters that model tonic and phasic firing of dopamine neurons. (D) Developmental stage did not impact levels of dopamine release in the DMS or (E) DLS. Insets: Individual data points for single-pulse dopamine release. In line graphs, symbols represent means ± SEMs. In bar graphs, bars represent means and symbols represent individual data points. *p < 0.05.

Dopamine release was initially evoked by a single electrical pulse (750 μA, 2 msec, monophasic) applied to the tissue every 3 minutes. Once the extracellular dopamine response was stable (3 collections within <10% variability), five-pulse stimulations were applied at varying burst frequencies (5, 10, 20, or 100 Hz) to model the physiological range of dopamine neuron firing. After assessing the dopamine response to single and multi-pulse stimulations, sulpiride (1μM), a dopamine D2/D3 receptor antagonist, was bath applied and dopamine response was equilibrated to single pulse stimulation and five-pulse stimulations were reassessed. On separate slices, single electrical pulse stimulations were applied until dopamine response was stable (as above). Once stable, quinpirole, a selective dopamine D2/D3 receptor agonist, was bath applied in increasing concentrations (3 nM - 1 μM), stabilizing dopamine response between each ascending dose.

### 2.4 Data Analysis

Demon Voltammetry and Analysis software was used to acquire and model FSCV data (Yorgason et al., 2011). Recording electrodes were calibrated by recording electrical current responses (in nA) to a known concentration of dopamine (3 μM) using a flow-injection system. This was used to convert electrical current to dopamine concentration. Michaelis-Menten kinetics were used to determine maximal rate of dopamine uptake (*V*max) (Ferris et al., 2013).

### 2.5 Statistics

Baseline single- and multi-pulse dopamine release and percent changes in dopamine release following sulpiride application were compared by two-factor ANOVA. In the case of significant interactions, Bonferroni or Tukey post-hoc comparisons were used. Dopamine release following quinpirole application was transformed into log scale and a non-linear regression was used to generate IC_50_ and area under the curve (AUC) for each curve. Mean IC_50_ were compared by extra sun-of-squares F test. Mean AUC and single pulse dopamine release were compared by students t-test. Graph Pad Prism (version 8, La Jolla, CA) or SPSS (version 24, International Business Machine Corporation, Armonk, NY) were used to statistically analyze data sets and compose graphs. Values >2 standard deviations above or below the mean were considered outliers and excluded. Data are presented as mean ± SEM.

## 3. Results

### 3.1 Decreased NAc dopamine release in adolescent rats

We first compared stimulated dopamine release in the striatum of adolescent (P31-37) and adult (>P70) male rats. To examine dopamine release at frequencies that model tonic- and phasic-like firing patterns, we stimulated single pulse and five pulses across the range of physiological dopamine firing rates. Figure 1A shows representative traces of single pulse stimulated dopamine release across four regions of the striatum. Developmental phase did not impact dopamine release in the NA shell, DMS, or DLS (Figure 1B, D, and E). However, adolescents had significantly less dopamine release in the NAc in response to the full range of stimulation frequencies (interaction: *F*_4, 40_=3.473, *p*=0.0158) and to single-pulse stimulations (*t*_10_=4.3, *p*=0.0016)(Figure 1C).

Activity of the dopamine transporter (DAT) can impact evoked dopamine release (see Ferris et al., 2013). To examine whether DAT uptake was mediating differences in dopamine release, we examined the maximal uptake rate of dopamine (*Vmax)* across the striatum (Figure 2A). There was no significant difference in *Vmax* between adolescents and adults in the NA shell, NAc, or DLS (Figure 2B, C, and E). However, *Vmax* was significantly greater (i.e. faster maximal dopamine uptake) in adolescents in the DMS (*t*_21_=2.212, *p*=0.0382)(Figure 2D).

**Figure 2.**
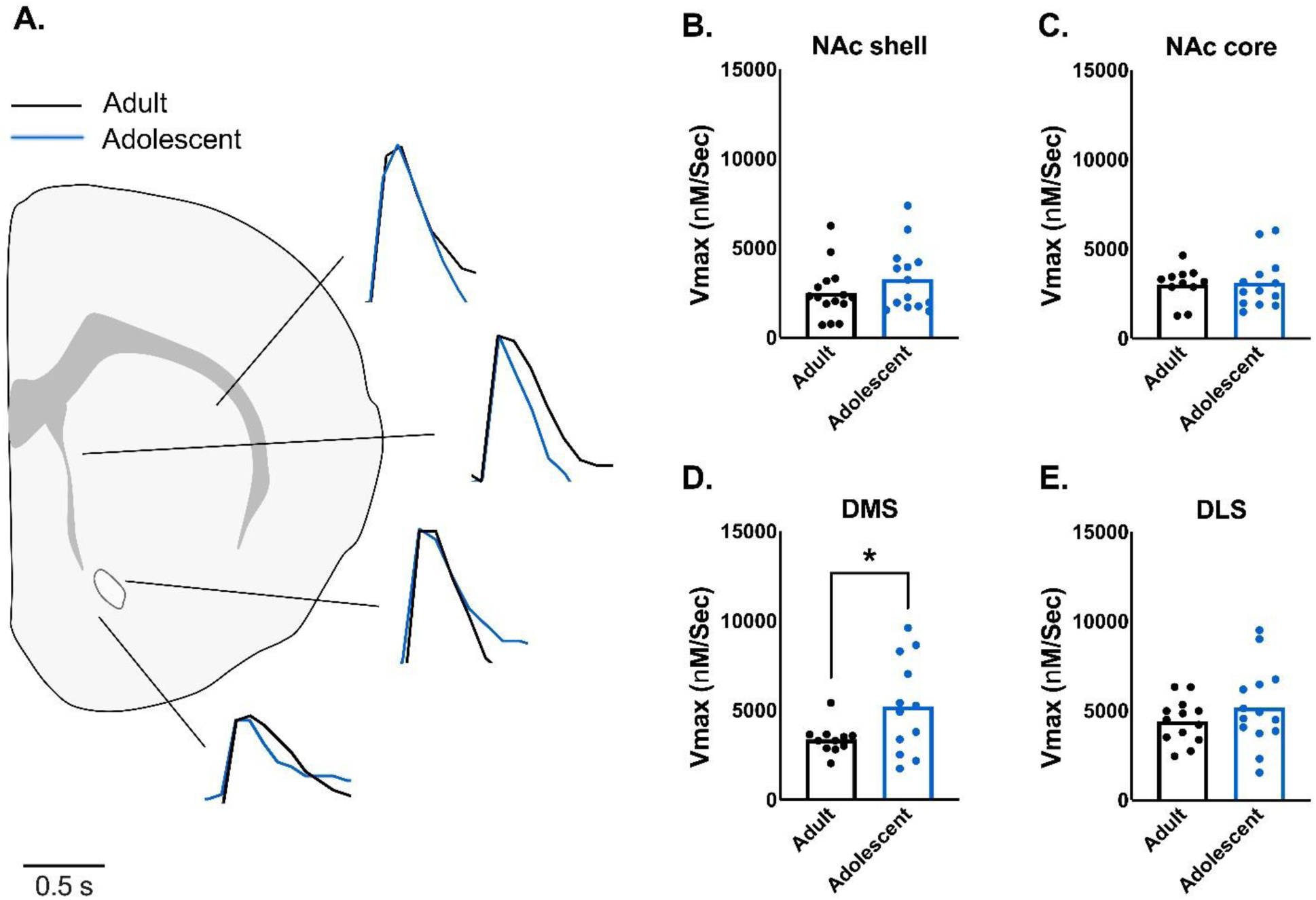
Maximal rate of dopamine uptake is faster in the dorsomedial striatum of adolescent rats. (A) Maximal rate of dopamine uptake (*Vmax*) was examined using *ex vivo* fast scan cyclic voltammetry in adult and adolescent rats. Representative traces of dopamine release and uptake in the nucleus accumbens shell (NA shell), nucleus accumbens core (NAc core), dorsomedial striatum (DMS), and dorsolateral striatum (DLS) are shown. Peaks of dopamine release were aligned to best compare rate of uptake. (B) Vmax was not different between adults and adolescents in the NA shell or (C) the NAc core. (D) Vmax was significantly higher in the DMS of adolescent rats, indicating faster uptake rates. (E) However, Vmax did not differ by age in the DLS. Bars represent means and symbols represent individual data points. *p < 0.05.

### 3.2 Increased D2/D3 receptor sensitivity in adolescent rats

Next, we hypothesized that autoreceptors may be mediating the developmental differences in stimulated dopamine release in the NAc, since D2 autoreceptors play a large role in mediating dopamine release (Zhang & Sulzer, 2012). Importantly, *ex vivo* FSCV is an excellent technique for isolating the contribution of autoreceptor on their respective neurotransmitter signals (Maina & Mathews, 2010; Bello et al., 2011). To examine this, we bath applied increasing doses of quinpirole, a selective D2/D3 receptor agonist, and calculated an inhibition dose effect curve for each striatal region (Figure 3A-D). The IC_50_ of the inhibition dose effect curve was not impacted by age for any region of the striatum (data not shown). However, the area under the curve (AUC) was significantly lower in the NAc of adolescents (*t*_8_=3.1, *p*=0.0147), but no different in any other striatal regions (Figures 3A-D insets).

**Figure 3.**
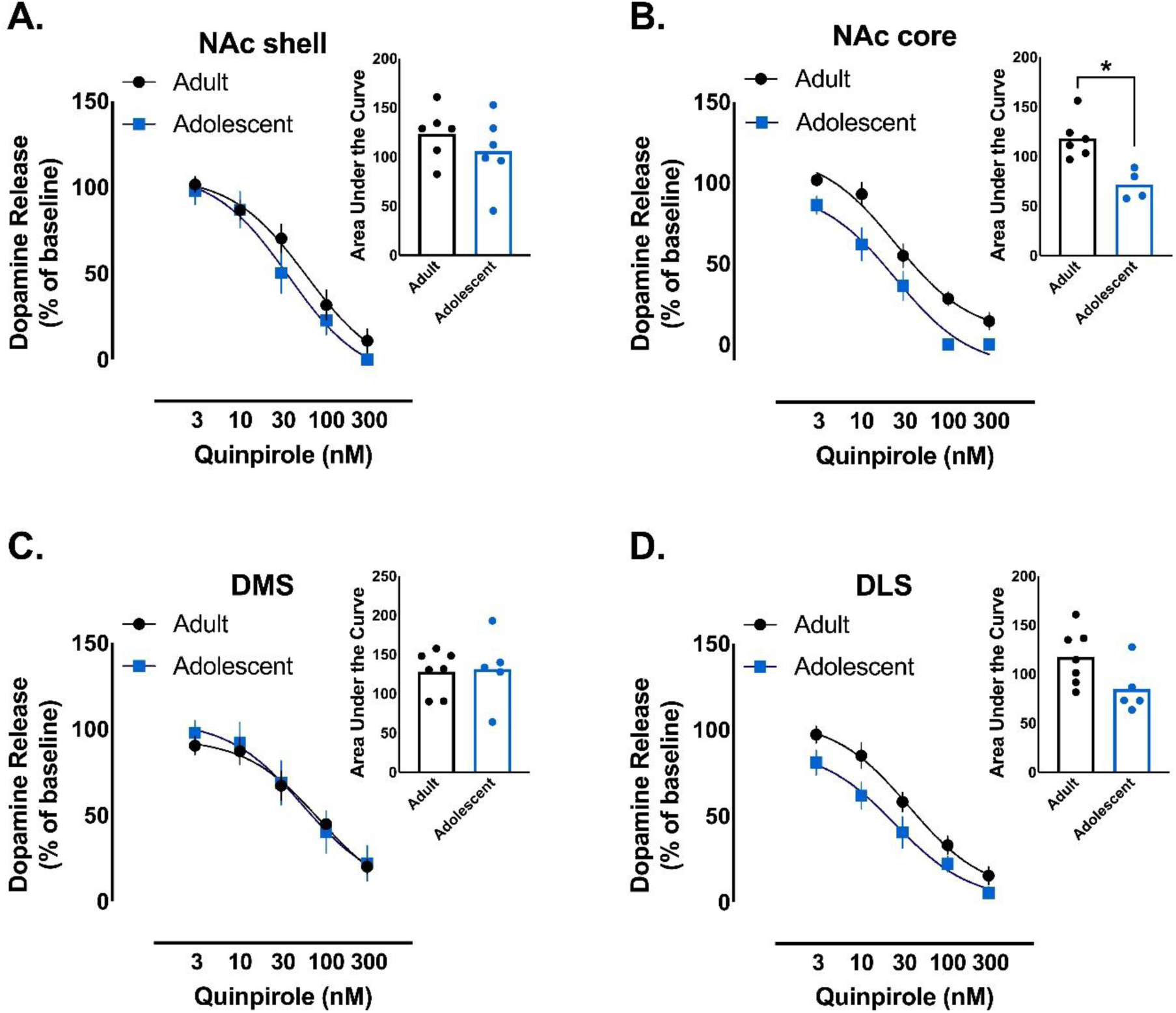
Dopamine autoreceptors in the nucleus accumbens core are hypersensitive in adolescent rats. (A) To examine the role of dopamine autoreceptors on dopamine release we bath applied increasing doses of quinpirole, a dopamine D2/D3 receptor agonist, and generated inhibition curves. Quinpirole decreased dopamine release in the nucleus accumbens shell (NA shell), but the inhibition curve was not differentially impacted by age. (B) In contrast, adolescents have a down-shifted quinpirole inhibition curve in the nucleus accumbens core (NAc core) and a significantly decreased area under of the curve (inset). This indicates adolescents have more sensitive dopamine autoreceptors than adults in the NAc core. Interestingly, the slope of the curve is not different between adolescents and adults, indicating there is not a potency shift in quinpirole’s effects. (C) Quinpirole inhibition curves were similar in adults and adolescents in the dorsolmedial striatum (DMS) and (D) dorsolateral striatum (DLS). Insets: Area under the curve values for each rat’s inhibition curve. In line graphs, symbols represent means ± SEMs. In bar graphs, bars represent means and symbols represent individual data points. *p < 0.05.

To further characterize striatal dopamine autoreceptors, we examined single and multi-pulse dopamine release following bath application of sulpiride, a D2/D3 receptor antagonist. We did not see any developmental differences in changes in dopamine release following sulpiride application in any examined striatal region (Figure 4A-D).

**Figure 4.**
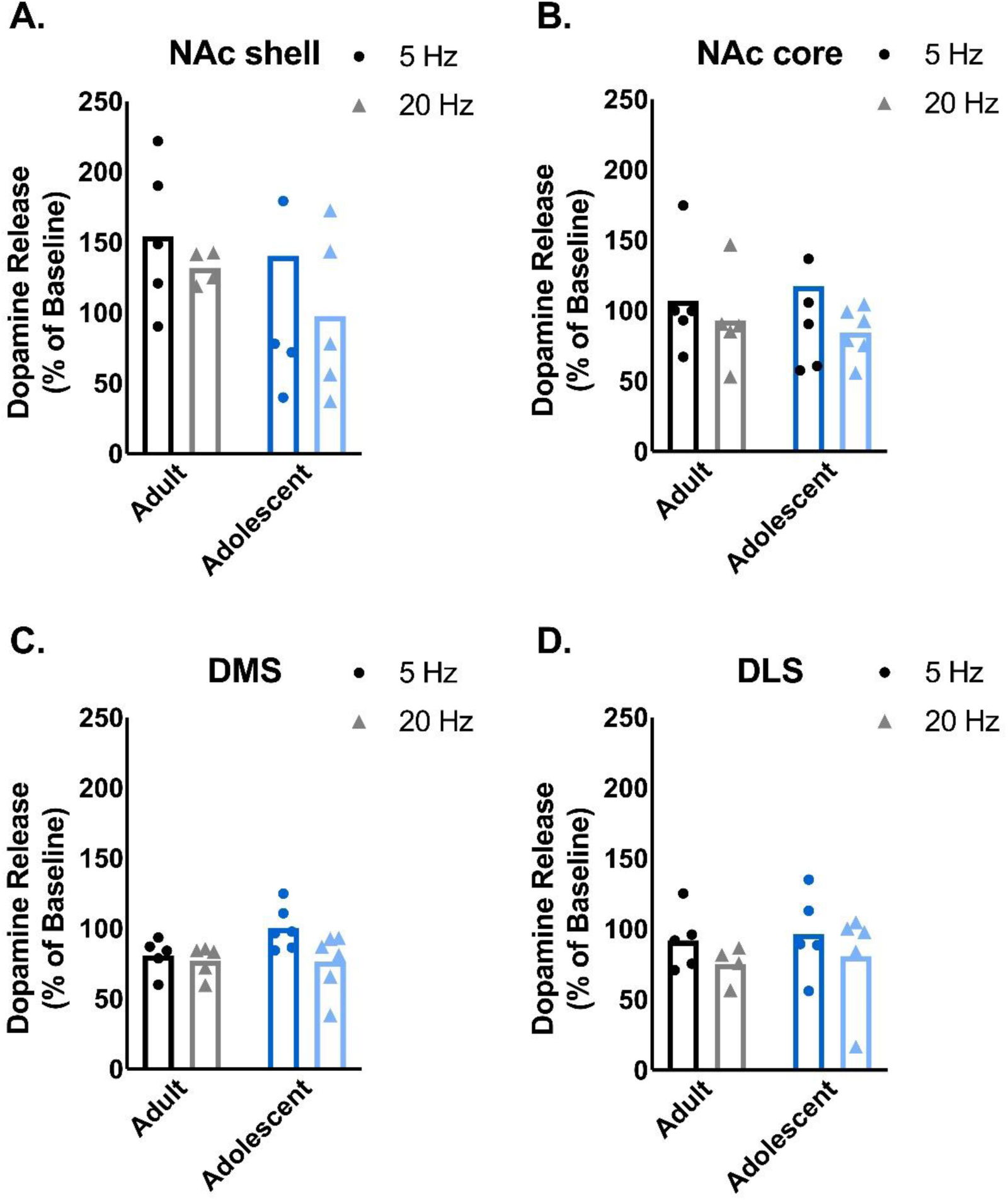
Sulpiride, a dopamine D2/D3 receptor antagonist, does not differentially impact dopamine release in adult and adolescent rats. (A) Sulpiride (1 μM) was bath applied to slices and the percent baseline dopamine release was compared across adult and adolescent animals for multi-pulse stimulations that model both tonic (5 Hz) and phasic (20 Hz) firing of dopamine neurons. Age did not impact the effect of sulpiride on dopamine release in the nucleus accumbens shell (NA shell), (B) nucleus accumbens core (NAc), (C) dorsomedial striatum (DMS), or (D) dorsolateral striatum (DLS). Bars represent means and symbols represent individual data points.

## 4. Discussion

Here, we utilized *ex vivo* FSCV to compare stimulated dopamine release and uptake in adolescents and adults across the striatum. Adolescence is characterized by increases in risk-taking, sensation-seeking, and peer-focused sociality (Spear, 2000, 2011). It is also a time of increased vulnerability to the development of a range of psychiatric disorders, including SUDs and mood and anxiety disorders (Casey et al., 2008; Fareri et al., 2008; Steinberg, 2008). The dopamine system has been shown to play an essential role in a number of psychiatric disorders, and the reorganization of the dopamine system during adolescence has been implicated in the unique behavioral changes and vulnerability of this developmental stage (Spear, 2013; Nelson et al., 2005; Wahlstrom et al., 2010a,b). Thus, our study aimed to characterize adolescent differences in striatal dopamine dynamics and its local circuitry regulation.

Our results indicate that adolescent rats have lower dopamine release than adults in the NAc core across a range of frequencies that model tonic and phasic firing of dopamine neurons. Interestingly, dopamine release was not impacted by developmental stage in other examined regions of the striatum. Differences in stimulated dopamine release in the NAc core could play an important role in unique changes in adolescent reward-related behaviors and vulnerability to psychiatric disorders. The NAc core plays an essential role in learning and reward-associated behaviors (Everitt & Robbins, 2005) and it is necessary for goal-directed decision making and incentive motivated behaviors (DiChiara, 2002; Saddoris et al., 2013). This positions dopamine release in the NAc core to play an important role in increased risk-taking in adolescents, which is hypothesized to be driven by increased incentive-reward motivation (Wahlstrom et al., 2010a). Additionally, it has also been hypothesized that the NAc core plays a role in the pathophysiology of mood disorders, including depression (Nestler & Carlezon, 2006). That adolescents, a more vulnerable population, has *decreased* dopamine release in the NAc core was unexpected. However, decreased dopamine release in the NAc core is seen in adults following chronic administration of a variety of drugs of abuse, including cocaine and alcohol, and this decrease is thought to play a role in mediating vulnerability to substance use disorders (Martinez et al., 2005; Addy et al., 2010; Ferris et al., 2012). That adolescents show naturally decreased dopamine release could impact their response to drugs of abuse and rewarding stimuli, increasing their vulnerability to a range of psychiatric disorders. Moreover, adolescents have increased D1 and D2 receptor density (Teicher et al., 1995; Tarazi et al., 1999; Philpot et al., 2009), so future studies are needed to understand how post-synaptic response to dopamine release is altered in adolescents compared to adults. There are obviously other developmental changes that occur in the brain during adolescence that also play a role in mediating behavioral shifts during adolescence (see Spear, 2000, 2013). However, these findings indicate there are significant differences in local circuitry modulation of terminal dopamine release in the NAc core of adolescent rats, which may be an important mediator of adolescent behavioral changes and vulnerability.

Next, we wanted to examine additional dopamine dynamics that could be mediating the decrease in stimulated dopamine release seen in the NAc core of adolescent rats. The activity of the dopamine transporter (DAT) can influence terminal dopamine release (see Ferris et al., 2013). However, we did not observe differences in the rate of maximal dopamine uptake (*Vmax*) in the NAc core, indicating that the DAT functions similarly in adolescents and adults. The only striatal region in which we found developmental differences in *Vmax* was the DMS, where we saw increased rate of dopamine uptake. Higher rates of dopamine uptake are likely driven by an increase in dopamine transporter levels (Salahpour et al., 2008; Ferris et al., 2013). DAT levels can directly influence behavior, with mutant mice with decreased DAT levels in the DMS showing impairments in reversal learning (Cybulska-Klosowicz et al., 2017). The DMS is necessary for goal-directed actions and flexible decision making (Balleine & O’Doherty, 2010; Hart et al., 2014) and, interestingly, adolescents show increases in flexible decision making on tasks that are mediated by the DMS (Johnson & Wilbrecht, 2011). Thus, increased DAT function in the DMS of adolescents may have functional implications for decision-making strategies.

Next, we hypothesized that stimulated dopamine release may be mediated by developmental differences in the sensitivity of autoreceptors. Autoreceptors can impact terminal dopamine release and *ex vivo* FSCV is an ideal technique for examining their role in mediating dopamine release (Maina & Mathews, 2010; Bello et al., 2011). We observed no developmental shift in potency (as indicated by IC_50_) of the effects of quinpirole, a selective D2/D3 dopamine receptor agonist, across the striatum. However, there was a downward curve shift and a significantly reduced AUC of the inhibitory curve in the NAc core of adolescents. This shift in efficacy indicates that adolescents have hypersensitive D2/D3 receptors in the NAc core, which may play a role in decreasing NAc core stimulated dopamine release.

Surprisingly, however, the D2/D3 antagonist sulpiride did not differentially impact dopamine release in any region of the striatum in adolescents. This seemingly counters the increased sensitivity we observed in the NAc core of adolescents following administration of a D2 agonist. One possible explanation for this difference between agonist and antagonist is that our dopamine stimulation parameters were chosen to model endogenous firing patterns of dopamine neurons. Natural stimulation parameters, however, may not isolate differences in D2 modulation of dopamine release, particularly because there is no dopamine tone that can constitutively activate D2 receptors in a slice preparation (Phillips et al., 2002). Moreover, our stimulation patterns (2 ms duration) likely did not evoke dopamine for enough time to activate D2 receptors, as previous work examining the impact of sulpiride on dopamine release found that sulpiride does not impact dopamine release unless the duration of stimulation exceeds 500 ms (Wieczorek & Kruk, 1995). Another possible interpretation is that the differential effects of quinpirole are being driven by quinpirole-induced inhibition of acetylcholine (Stoof & Kebabian, 1982), a neurotransmitter known to regulate dopamine terminal release (Rice & Cragg, 2004). Acetylcholine is tonically released in the striatum, and pauses in acetylcholine signaling drive changes in striatal dopamine release (Cragg, 2006). Since striatal dopamine release is modulated through decreases in acetylcholine levels, and acetylcholine is degraded rapidly at the synapse, the inhibition in acetylcholine release caused by quinpirole may alter dopamine release more than any increases caused by sulpiride application. This interpretation would also indicate that acetylcholine signaling differentially regulates dopamine release in the nucleus accumbens of adolescent rats and future studies could examine this possibility.

To summarize, adolescent rats have decreased stimulated dopamine release in the NAc core across a range of tonic- and phasic-like firing patterns. This difference is not driven by changes in DAT functionality, but increased sensitivity of dopamine autoreceptors may play a role in decreasing dopamine release in the NAc core of adolescents. However, further studies are needed to confirm this hypothesis. Additionally, adolescents exhibit increased maximal rates of dopamine uptake in the DMS. We did not observe developmental differences in dopamine release or dynamics in the NA shell or DLS. These differences in dopamine release and their regulation may play a role in the unique behavioral changes and increased vulnerability unique to the adolescent developmental period.

